# Reanalysis of mtDNA mutations of human primordial germ cells (PGCs) reveals NUMT contamination and suggests that selection in PGCs may be positive

**DOI:** 10.1101/2022.12.12.520138

**Authors:** Zoë Fleischmann, Auden Cote-L’Heureux, Melissa Franco, Sergey Oreshkov, Sofia Annis, Mark Khrapko, Dylan Aidlen, Konstantin Popadin, Dori C. Woods, Jonathan L. Tilly, Konstantin Khrapko

**Affiliations:** Department of Biology, Northeastern University, Boston, Massachusetts, USA; School of Life Sciences, École Polytechnique Fédérale de Lausanne, 1015 Lausanne, Switzerland; Swiss Institute of Bioinformatics, 1015 Lausanne, Switzerland; Center for Mitochondrial Functional Genomics, Institute of Living Systems, Immanuel Kant Baltic Federal University, 236040 Kaliningrad, Russia

**Author notes:** equal contribution (Zoë Fleischmann, Auden Cote-L’Heureux, Melissa Franco).

## Abstract

The resilience of the mitochondrial genome (mtDNA) to a high mutational pressure depends, in part, on negative purifying selection in the germline. A paradigm in the field has been that such selection, at least in part, takes place in primordial germ cells (PGCs). Specifically, Floros et al. (*Nature Cell Biology* 20: 144–51) reported an increase in the synonymity of mtDNA mutations (a sign of purifying selection) between pooled early-stage and late-stage PGCs. We re-analyzed Floros’ et al. pooled PGC data and noticed that their mutational dataset was significantly contaminated with single nucleotide variants (SNVs) derived from a nuclear sequence of mtDNA origin (NUMT) located on chromosome 5. Contamination was caused by co-amplification of the NUMT sequence by cross-specific PCR primers. Importantly, *when we removed NUMT-derived SNVs, the evidence of purifying selection was abolished*. In addition to pooled PGCs, Floros et al. reported the analysis of *single* late-stage PGCs, which were amplified with different sets of PCR primers that cannot amplify the NUMT sequence. Accordingly, we found no NUMT-derived SNVs among single PGCs mutations. Interestingly, single PGC mutations show a *decrease* of synonymity with increased intracellular mutant fraction. This pattern is incompatible with predominantly negative selection. This suggests that germline selection of mtDNA mutations is a complex phenomenon and that the part of this process that takes place in PGCs may be predominantly positive. However counterintuitive, positive germline selection of detrimental mtDNA mutations has been reported previously and potentially may be evolutionarily advantageous.

## Introduction

Mitochondrial DNA (mtDNA) is known for having a high mutational rate and a high density of crucial genes. Therefore, mtDNA mutations not only underlie a spectrum of mitochondrial diseases (Lander and Lodish 1990), (Elliott et al. 2008) but also, if left unpurged, could result in long-term detrimental effects on species evolution, a.k.a. Muller’s ratchet (Popadin et al. 2007). It is thus vital to understand how mtDNA mutations are kept at a sustainable level. It has been long realized that detrimental mtDNA mutations are selected against not only at the level of the organism but also at the level of germ cells. This phenomenon was demonstrated in a 2008 study (Fan et al. 2008), where an efficient selection against highly detrimental mtDNA mutation was demonstrated in the mouse germline prior to ovulation. Another study from 2008 demonstrated significant levels of purifying selection against a wide range of detrimental mutations in the germline. This conclusion was based on the high synonymity of mtDNA mutations as early as the second generation after they were generated *de novo* in an mtDNA ‘mutator’ mouse line (Stewart et al. 2008). Similarly, purifying selection was reported to shape human mtDNA diversity (Wei et al. 2019). However, the mechanism(s) and precise developmental timing(s) of germline purifying selection are still under investigation.

Recently, Floros and colleagues (Floros et al. 2018) studied changes in the transcriptome and mtDNA mutations during the late-stage development of human primordial germ cells (PGCs). Importantly Floros et al. reported a sharp increase in synonymity of mtDNA mutations in late-stage (CS20/21: Carnegie Stage 20/21) PGCs, compared to early-stage (CS12), which they interpreted as purifying selection in late PGCs. However, the observed increase in synonymity of mutations in pooled late-stage PGCs appeared to result, surprisingly, *not from a decrease* of nonsynonymous mtDNA mutations as would be expected under negative purifying selection, but instead mostly from an *increase* in the frequency of synonymous mutations. In search of an explanation, we analyzed Floros et al. primary *pooled* PGC data as well as *single* PGC data, which were provided to us by Dr. Chinnery and are now published (Floros et al. 2022). We found that the apparent synonymity shift in PGCs resulted from NUMT co-amplification, a previously described artifact (Khrapko et al. 1994), (Hirano et al. 1997), (Wallace et al. 1997), (Lutz-Bonengel and Parson 2019) (Annis et al. 2019), (Balciuniene and Balciunas 2019), (Wei et al. 2020). Removal of NUMT-derived SNVs abolished evidence of purifying selection; moreover, analysis of non-contaminated *single* PGC data implied predominantly positive selection in these cells.

## Results and discussion

An initial review of Floros et al. primary data of *pooled* PGCs revealed that a majority (∼80%) of ‘approved’ sequence variants (i.e., those above the 1% threshold and thus deemed true mutations by the authors) were found in multiple unrelated samples (**Suppl. Table**, orange shading). Such profuse recurrence is normally not observed in somatic mutations, most of which are individually rare events and therefore are negligibly likely to occur in multiple samples. One widely recognized cause of such inter-sample repetition of low fraction mtDNA sequence variants is contamination with nuclear DNA of mitochondrial origin (NUMTs).

NUMT contamination is a well-recognized problem of mtDNA mutational analysis, which we and others first encountered decades ago (Khrapko et al. 1994), (Hirano et al. 1997). NUMTs are nuclear pseudogenes derived from fragments of mtDNA inserted into the nuclear genome. Many of them were inserted millions of years ago and since then nuclear and mitochondrial sequences have been diverging from each other. A typical NUMT differs from mtDNA by multiple changes, which appear as multiple linked SNVs and constitute a ‘haplotype’ specific to each NUMT. We tested the NUMT contamination hypothesis and were able to prove it with three lines of evidence.

First, we aligned the Floros et al. sequence variants to several known NUMT sequences of the human genome. This analysis revealed that at least ***30%*** of protein-coding single-nucleotide variants (SNVs) reported by Floros et al. as real mutations, actually mapped to a NUMT located on chromosome 5 (CNVs marked ‘ch5’ in **Suppl. Table**). Coincidentally, we have previously used this very NUMT as a marker of human evolution (Popadin et al. 2022) so we were very familiar with it. We determined that contamination was introduced by co-amplification of the NUMT sequences by one of the PCR primer pairs used in their study. Indeed, all NUMT-mapping SNVs were located in the mtDNA sequence between the primers of a primer pair that was used by Floros et al. to amplify one of their PCR fragments (**Fig. 1**, see also **Suppl. Note 1**).

**Figure 1.**
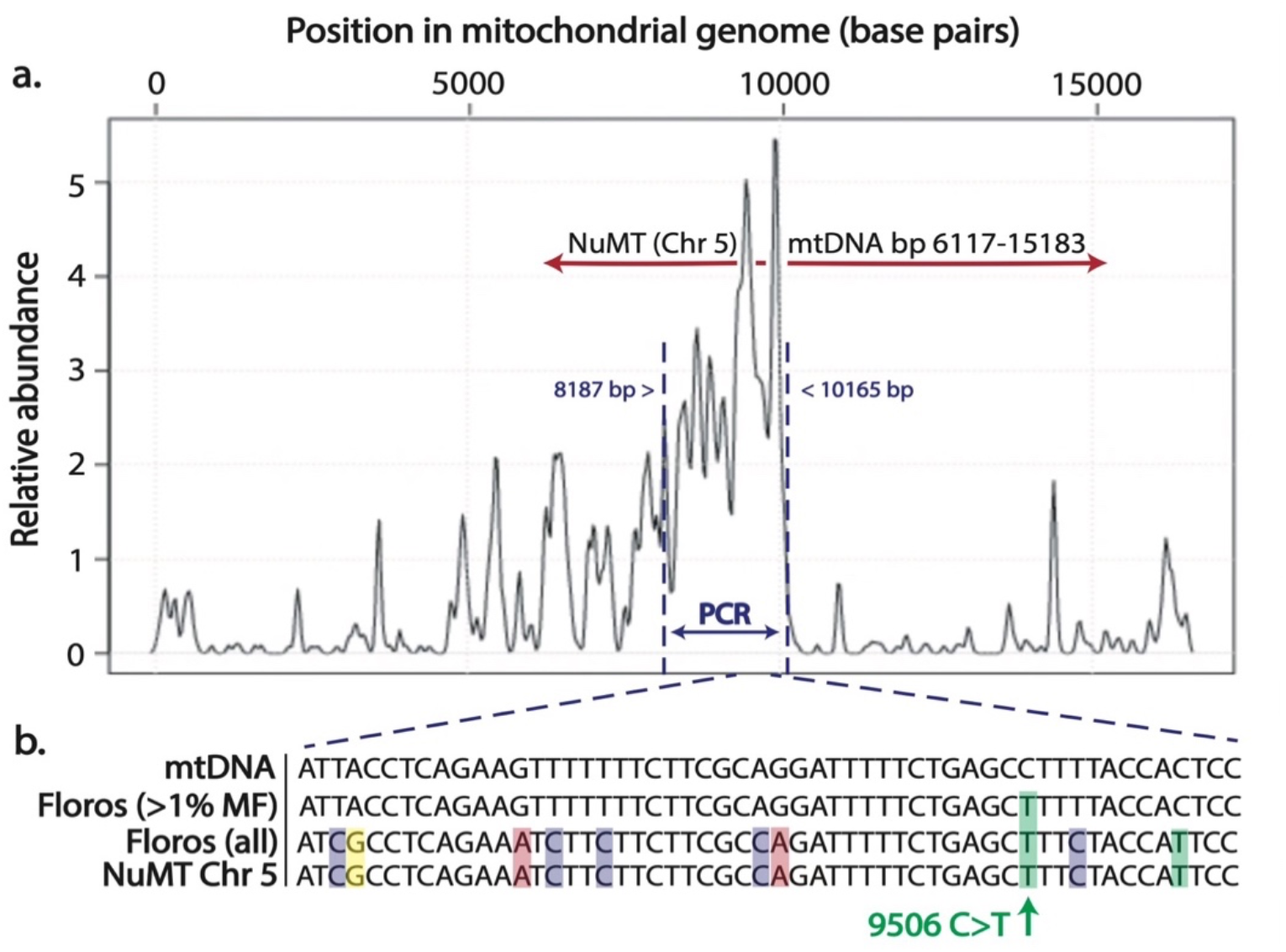
NUMT co-amplification contaminates mtDNA mutations from pooled PGCs: evidence from spatial distribution and sequence alignment. **A**. Spatial density distribution of SNVs from pooled PGC dataset (Floros et al 2018). The unexpected dense patch of variants (between ∼8kb and 10Kb, marked by dashed vertical lines) corresponds to one of the PCR fragments used in Floros et al. (bp:8187-10165, horizontal blue double arrow with the ‘PCR’ label). The PCR primers of this fragment co-amplify a portion of the nuclear pseudogene of mtDNA (NUMT) on Chromosome 5 (hg38(100045938-100055045)) that is homologous to this region of mtDNA (red double arrow; see also **Suppl. Note 1**). **B**. A representative section of alignment of the reference mtDNA sequence (‘mtDNA’) with the NUMT sequence on Chromosome 5 (‘NuMT Chr 5’) and two mtDNA sequences with added SNVs reported by Floros et al. Sequence “Floros (all)” includes all reported SNVs. “Floros (>1% MF)” includes only the ‘approved’ SNVs that are above 1% threshold. This alignment shows that a great majority of SNVs from Floros et al. are identical to the NUMT-derived SNVs, consistent with their NUMT origin. Of 10 variants located in this section of the genome, one (9506C>T, green arrow) exceeded the threshold and thus was erroneously considered a real mutation. 9506C>T is just one example; there are 11 cases like that across this PCR fragment (see **Suppl. Table**). This contamination is substantial in relative terms: NUMT-derived SNVs account for 10 of 28 protein-coding SNVs in late-stage PGCs (36%) and 1 of 8 in early PGCs (12%). See **Fig.S1** for more details.

Second, NUMT origin of NUMT-mapping SNVs was confirmed by comparison to pooled PGC mutations to single PGC mutations. Floros et al amplified Single PGCs using different sets of primers than pooled PGCs. We determined that none of the two primer pairs used for single PGCs amplification were able to amplify any portion of this NUMT (shown in schematics in **Suppl. Note 2)**. In full agreement with NUMT co-amplification hypothesis, no NUMT - mapping SNVs were detected in single PGC samples.

Third, the strongest evidence of NUMT origin of NUMT-mapping SNVs was obtained using raw next-generation sequencing (NGS) data (provided by Floros et al in their Supplement). We determined that NGS reads of the pooled PGCs fell into two classes – the ‘mtDNA-derived reads’ and the ‘NUMT-derived reads’. The latter were clearly distinguished by the presence of the ‘NUMT haplotype’, i.e., a full set of NUMT-specific nucleotide changes present in a single NGS read. Note that *all* 1%+ NUMT-mapping SNVs reported by Floros et al. were derived exclusively from the NUMT-derived NGS reads, which carry the full NUMT haplotype, definitively proving their NUMT origin (see Fig. S1 and its caption for details of the analysis).

Having confirmed that NUMT-mapping SNVs are indeed NUMT-derived, we asked whether contamination with NUMT-derived SNVs affected the key conclusions of the study and in which way. The effect of the removal of NUMT contamination on the estimated synonymity change between early and late PGCs is shown in **Figure 2**. In this figure, we recreated original data plots from Figures 2b and 2c of the Floros study (these are left-side panels 2a and 2c of *our* **Figure 2**, correspondingly). Then we removed NUMT-derived SNVs and re-plotted the data (right side panels Figure 2b and d, correspondingly).

**Figure 2.**
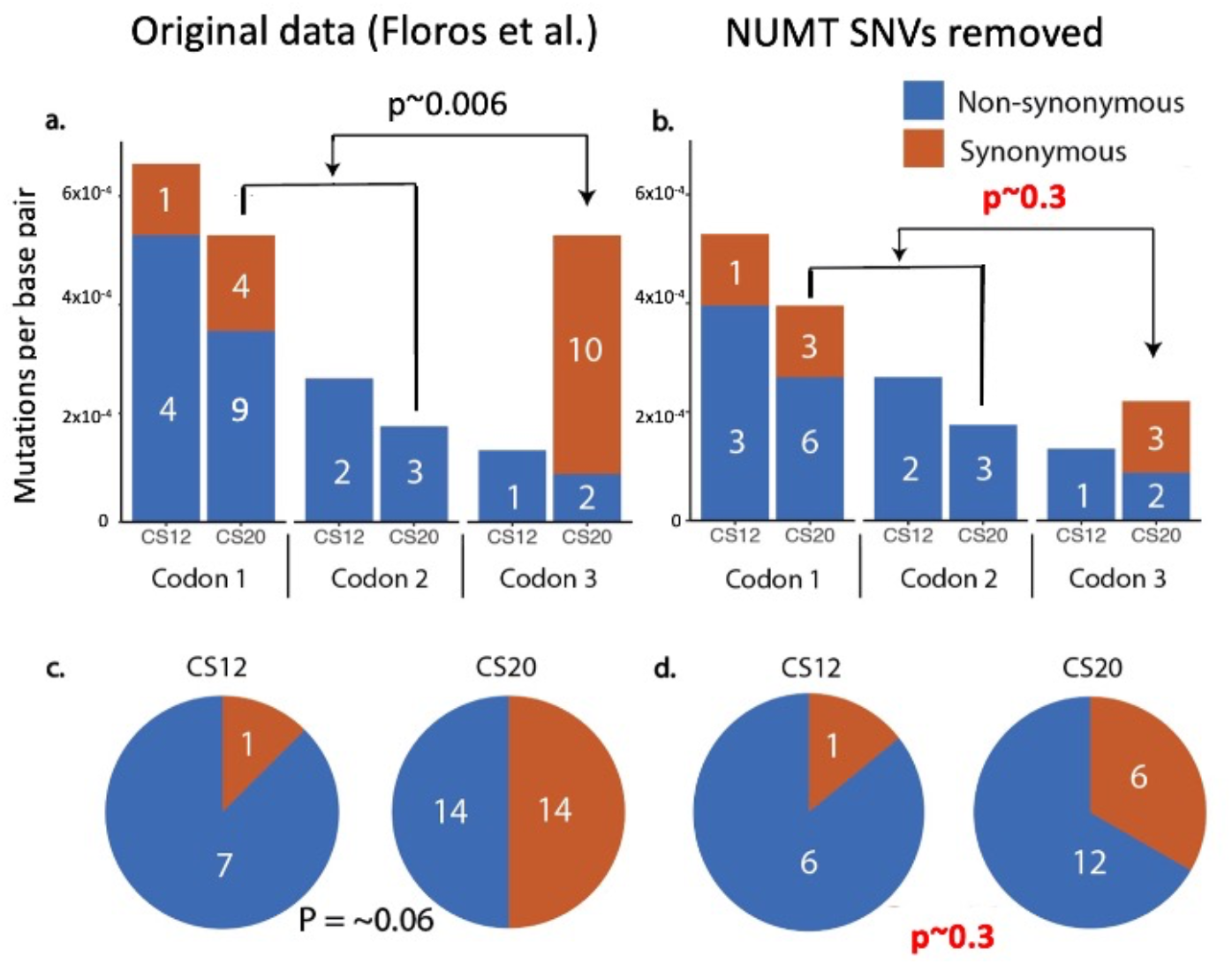
Removal of NUMT contamination fully abolishes statistical evidence of purifying selection in late-stage PGCs. **a:** a recreation of Figure 2b from Floros et al., 2018. **b:** The same graph after the removal of NUMT-mapping SNVs. **c:** A recreation of Figure 2c from Floros et al., 2018 using the original data and **d:** pie charts after the removal of NUMT-derived SNVs. We note that the removal of NUMT contamination illustrated in this figure is very conservative. P-values were calculated using Fisher Exact Test. White numbers within bars and pie charts represent the absolute number of mutations in each category; note that these numbers are not proportional to the size of the bars because the mutant fraction represented by the height of the bars is calculated relative to the number of embryos in each category, and there are 3 times more late stage embryos (6) than early stage embryos (2).

As seen in **Figure 2**, the removal of NUMT-derived SNVs results in a *complete loss* of statistical significance of the synonymity change between early and late-stage PGCs both in overall synonymity (p-value increases from 0.06 with NUMTs to 0.34 without NUMTs, Fig.2 c vs. d) and in codon bias (p-value increases from 0.006 with NUMTs to 0.3 without NUMTs, **Fig.2a vs. b**). In conclusion, the apparent synonymity change and the purifying selection deduced from this change are merely a result of NUMT contamination.

We then asked *how* NUMT contamination caused such inaccurate assessment of the synonymity change between early and late PGCs. **Supplementary Table** shows that the NUMT contamination is biased: 10 of NUMT-derived SNVs are in late-stage PGC samples, and only one is in early-stage PGCs. This is a 3-fold bias, after correction for the number of mutations. Note that NUMT-derived SNVs are generally highly synonymous. This is because, due to a much higher mutational rate in mtDNA than in nuclear DNA, most differences between a NUMT and mtDNA (which are recorded as NUMT SNVs) are in fact ancient mtDNA mutations, absolute majority of which which are synonymous (this is further discussed in **Suppl. Note 3**, see also (Popadin et al. 2022)). Thus, an excess of synonymous, NUMT-derived SNVs in late PGC naturally led to a significant overestimation of the synonymity of their mutations, and therefore to the perceived purifying selection in late PGCs. Reassuringly, this prevalence of synonymous NuMT-derived SNPs fully explains the puzzling surge of synonymous variants in late-stage PGCs (**Fig.2a**) which initiated this inquiry.

Why is NUMT contamination in Floros et al. so biased towards late PGCs? **Suppl. Table** shows that almost all NUMT-derived SNVs (8 out of 10) come from a single late-stage PGC sample, CS20-11593. For an explanation of this peculiarity, we turned to unprocessed mutational data (Floros et al., Table S5). This table contains all SNVs detected in pooled PGC samples, including those below the 1% threshold, down to 0.3%. We found that NUMT-derived SNVs were in fact detected in all pooled PGC samples, however, in the CS20-11593 sample a large proportion of these SNVs exceeded 1%, whereas in other samples most of them remained just below 1%. Even in sample CS20-11593 most of the NUMT-derived SNVs were ‘hidden’ just under the 1% threshold. The variability of mutant fraction between different NUMT-derived SNVs may look unexpected because, as we noted before, all NUMT-derived SNVs are part of a haplotype and are linked to each other on sequencing reads. The likely cause of this variability is discussed in **Supplementary figure S1**.

In addition to protein-coding mutations, Floros et al. reported a decrease in RNA-coding mutations in late PGC with a p-value of 0.03 (Figure 2d of Floros et al). We were not able to reproduce this result using Floros et al. data: there is only one non-coding RNA mutation in CS12 and only two in CS20/21. Finally, Floros et al. reported convincing evidence of selection among mutations of the control region. We note, however, that D-loop mutations, although many of them are known to be prone to selection, are typically not detrimental, making the status of D-loop mutations with respect to purifying selection unclear.

Because *pooled* PGC data proved to be contaminated with NUMTs and are inconclusive, we turned to *single* PGC data, which, as discussed earlier, are free from contamination with NUMT on ch5. The single PGC dataset is comprised of late-stage PGCs only, so early vs. late-stage comparison is not possible. We note, however, the single PGC data contain a large number of clonally expanded mutations (Floros et al., 2018), which permits an alternative way to assess purifying selection. Logically, purifying selection of an mtDNA mutation can proceed within or between cells. Within cells, purifying selection may preferentially remove mitochondria with detrimental mutations or prevent them from proliferation. Either way, clonal expansion of detrimental mutations is blocked. On the cellular level, selection may preferentially remove or prevent the proliferation of cells with a higher mutant fraction (i.e., larger clone size) of detrimental mutations. Either scenario predicts an increased proportion of synonymous mutations among mutations with higher intracellular mutant fractions, than those with lower MF. In other words, mutations with higher intracellular mutant fractions should have increased synonymity. Similarly, this logic predicts that average mutant fractions of non-synonymous mutations should be lower than average mutant fractions of synonymous ones.

With those expectations in mind, we tested the purifying selection hypothesis by testing the predictions. First, we noted that, in direct disagreement with the predictions, the average mutant fraction of *synonymous* single PGC mutations was significantly lower than that of *nonsynonymous* ones (0.028 vs. 0.042, *p<0*.*02*, two-sample t-test. The data used in these and following calculations can be found in suppl. Note 4). This effect is echoed by a dramatic decrease in synonymity at higher mutant fractions shown in **Figure 3**. Moreover, detailed analysis revealed an even stronger disagreement with the purifying selection hypothesis: we found that the difference in average mutant fraction between synonymous and nonsynonymous mutations was due to a group of statistical outliers, i.e., 12 nonsynonymous mutations with the highest mutant fractions in the entire dataset (see Suppl. Note 4 for details).

**Figure 3.**
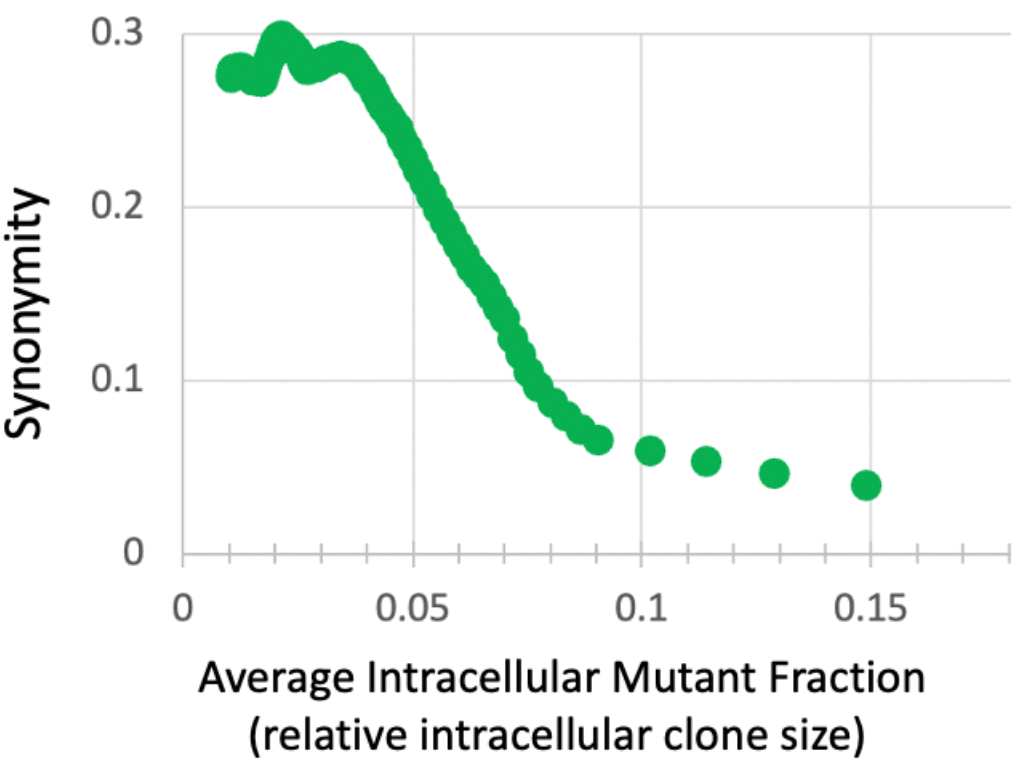
The synonymity of clonally expanded single PGC mutations decreases with the relative intracellular clone size (which is equivalent to the intracellular mutant fraction). To make this graph, single PGC mutations were ranked by their mutant fraction, then synonymity and average mutant fraction was calculated in sliding windows 20 mutations wide. The observed decrease of synonymity with average clone size is the opposite of what is predicted by the purifying selection hypothesis, moreover, it implies a positive selection of non-synonymous mutations in PGCs (see text for details). The statistical significance of this decrease is formally demonstrated in Supplemental note 4.

The prevalence of high-fraction nonsynonymous mutations with mutant fractions exceeding those of synonymous mutations strongly contradicts purifying selection either on the mitochondrion or the cellular level. However, the formal proof of the statistical significance of the observation of these high-fraction nonsynonymous mutations requires non-parametrical analysis (outliers, by definition, are not normally distributed). The binomial probability of observing this set of nonsynonymous outliers is 0.017, i.e., this observation is statistically significant with p<0.017. In addition, we performed bootstrap simulation which demonstrated that the non-parametric p-value of the enrichment of nonsynonymous mutations at high mutant fractions is highly significant (p∼0.01; see Suppl. Note 4 for details).

The latter findings imply that the hypothesis of predominant purifying selection in late PGCs is not merely ‘not supported’ by the data, but has been rejected. Moreover, a statistically significant increase of non-synonymity among clonally expanded mutations in individual PGCs implies predominantly *positive* selection in PGCs. We use the word ‘*predominant*’ to acknowledge that synonymity change only attests to the predominant direction of selection. The overall decrease in synonymity does not exclude that some detrimental mutations may be under purifying selection. All this means is that a predominant majority of nonsynonymous mutations are under positive selection. As a note of caution, this result pertains only to mutant fractions or 5% and higher, because below this level significance disappears, and it is possible that at lower mutant fractions there is no positive selection or selection becomes predominantly purifying. This conclusion of predominantly positive selection is quite extraordinary, however, given the relatively small size of the study it should be considered preliminary and needs independent confirmation.

As counterintuitive as it may seem, positive germline selection of detrimental mtDNA mutations is not a novel observation. An average increase of heteroplasmy (the term ‘heteroplasmy’ traditionally is used in place of mutant fraction when discussing inherited mutations) across generations has been reported for 8993T>G (Otten et al. 2018). More recently, it became clear that some mtDNA mutations may be subject to positive germline selection within a limited interval of heteroplasmy level. We have recently reported that two fairly detrimental mutations, mouse 3875delC and human 3243A>G, are subject to such fraction-dependent positive selection (Franco et al. 2020) and (Fleischmann et al. 2021). More recently, Chinnery’s laboratory also proposed positive selection at low mother’s heteroplasmy in the mouse 5024C>T and human 3243A>G, 8344A>G, and 8993 T>G (Zhang et al. 2021). Thus, positive selection of detrimental mutations in the germline may be a fairly common phenomenon.

How can the putative positive selection of detrimental mtDNA mutations be compatible with overall purifying selection in the germline, e.g., as reported by (Stewart et al. 2008) and (Wei et al. 2019)? We note that the signature of positive selection inferred in our study pertains to mutations with intermediate mutant fractions and happens during a specific interval of germline development. The significance ceases at lower mutant fractions (see Supplementary Note 4). This is consistent with our previous findings of an ‘arching’ selection profile in human m.3243A>G mutation (Franco et al. 2022). More generally, mutations at other mutant fractions, mutations of other types (e.g., highly detrimental mutations - (Fan et al. 2008)), and at different stages of germline development may be subject to purifying selection so that overall balance of selection in the germline is negative despite some positive intervals. This study confirms the potential complexity of the inner works of purifying germline selection. A recent report by the Chinnery group that a detrimental mtDNA mutation in the mouse (*m*.*5024C>T)* is subject to positive selection specifically during oocyte maturation (Zhang et al. 2021) is consistent with the idea of the complexity of germline selection. Interestingly, the site of positive selection that they report, is different from the one described here. More research is needed to reconcile these findings. The differences may be attributable to the special properties of the m.5024C>T mutation. m.5024C>T is known for its unusual inheritance pattern: for example, it was not possible to obtain heteroplasmy below 15% in the offspring (*Jim Stewart, personal communication*).

In conclusion, the pattern of germline selection appears to be potentially complex and dependent on the type of mutation, it’s a fraction in the cell and on the stage of germline development. More research is needed to explore this critical phenomenon and in doing so great caution should be exercised to avoid dangerous artifacts inherent to mutation analysis of mtDNA, including NUMT contamination.

## Methods

### Synonymity and repeatability analysis of sequence variants of mtDNA

The data of the Floros et al. study have been imported from their supplemental table 5 at https://doi.org/10.1038/s41556-017-0017-8 (Floros et al., 2018). Synonymity was analyzed using the ‘lookup’ function on in-house Excel-based spreadsheets containing synonymity tables for all possible mutations in the mitochondrial genome. Identical mutations in different samples were identified using the ‘countif’ function.

### Extraction and multiple alignment of raw reads from the Next Generation Sequencing (NGS) dataset

We extracted raw read NGS data from NCBI Sequence Read Archive record SRR6288291, which corresponds to the late PGC sample CS20-11593. We aligned reads against an mtDNA reference sequence (bp 9379-9436) that corresponds to the Chr5 NuMT with the BWA-MEM tool (http://bio-bwa.sourceforge.net/) and then filtered the reads to only capture those with seven or fewer mismatches using samtools (“[NM]<=7”) and had a mapping quality of ≥ 20; duplicate sequences were removed using a custom script. We again used samtools to convert .bam files to. fasta files.

***Multiple alignment*** *of the extracted reads* was performed using the NCBI BLAST server with default parameters, with the sequence of the human NuMT at chromosome 5 (hg38(ch5100045938-100055045)) as a query. We chose this method merely as a visually convenient approach to present the results.

## Abbreviations

CS: Carnegie Stage, (conventional staging of human embryonic development) followed by the stage number
PGC: Primordial Germ Cell
NUMT: a pseudogene of a mitochondrial DNA residing in the nuclear genome
MF: mutant fraction
SNV: single nucleotide variant. Used instead of ‘mutation’, especially where it is not clear whether the mutation is real or artificial

## Acknowledgments

This study was supported by a grant from the U.S. National Institutes of Health (R01-HD091439 to J.L.T., D.C.W. and K.K.), KP was supported by the Ministry of Science and Higher Education of the Russian Federation (agreement no. 075-02-2022-872). We are thankful to Dr. Chinnery for sharing his data prior to publication.

## References

Annis, Sofia, Zoe Fleischmann, Mark Khrapko, Melissa Franco, Kevin Wasko, dori woods, Wolfram S. Kunz, Peter Ellis, and Konstantin Khrapko. 2019. “Quasi-Mendelian Paternal Inheritance of Mitochondrial DNA: A Notorious Artifact, or Anticipated Behavior?” Proc Natl Acad Sci U S A 116 (30): 14797–98. https://doi.org/10.1073/pnas.1821436116.

Balciuniene, Jorune, and Darius Balciunas. 2019. “A Nuclear MtDNA Concatemer (Mega-NUMT) Could Mimic Paternal Inheritance of Mitochondrial Genome.” Frontiers in Genetics 10 (June): 518. https://doi.org/10.3389/fgene.2019.00518.

Elliott, H. R., D. C. Samuels, J. A. Eden, C. L. Relton, and P. F. Chinnery. 2008. “Pathogenic Mitochondrial DNA Mutations Are Common in the General Population.” Am J Hum Genet 83 (2): 254–60.

Fan, W., K. G. Waymire, N. Narula, P. Li, C. Rocher, P. E. Coskun, M. A. Vannan, J. Narula, G. R. MacGregor, and D. C. Wallace. 2008. “A Mouse Model of Mitochondrial Disease Reveals Germline Selection against Severe MtDNA Mutations.” Science 319 (5865): 958–62.

Fleischmann, Zoe, Sarah J. Pickett, Melissa Franco, Dylan Aidlen, Mark Khrapko, David Stein, Natasha Markuzon, et al. 2021. “Bi-Phasic Dynamics of the Mitochondrial DNA Mutation m.3243A>G in Blood: An Unbiased, Mutation Level-Dependent Model Implies Positive Selection in the Germline.” BioRxiv, January, 2021.02.26.433045. https://doi.org/10.1101/2021.02.26.433045.

Floros, Vasileios I., Angela Pyle, Sabine Dietmann, Wei Wei, Walfred C. W. Tang, Naoko Irie, Brendan Payne, et al. 2018. “Segregation of Mitochondrial DNA Heteroplasmy through a Developmental Genetic Bottleneck in Human Embryos.” Nature Cell Biology 20 (2): 144–51.

Floros, Vasileios I., Angela Pyle, Sabine Dietmann, Wei Wei, Walfred W. C. Tang, Naoko Irie, Brendan Payne, et al. 2022. “Author Correction: Segregation of Mitochondrial DNA Heteroplasmy through a Developmental Genetic Bottleneck in Human Embryos.” Nature Cell Biology, December. https://doi.org/10.1038/s41556-022-01046-z.

Franco, Melissa, Sarah Pickett, Zoe Fleischmann, Mark Khrapko, Sofia Annis, dori woods, Natalya Markuzon, Doug Turnbull, and Konstantin Khrapko. 2020. “Can Detrimental MtDNA Mutations Be under Positive Selection in the Germline?” The FASEB Journal 34 (S1): 1–1. https://doi.org/10.1096/fasebj.2020.34.s1.09461.

Franco, Melissa, Sarah J Pickett, Zoe Fleischmann, Mark Khrapko, Auden Cote-L’Heureux, Dylan Aidlen, David Stein, et al. 2022. “Dynamics of the Most Common Pathogenic MtDNA Variant m.3243A>G Demonstrate Frequency-Dependency in Blood and Positive Selection in the Germline.” Human Molecular Genetics 31 (23): 4075–86. https://doi.org/10.1093/hmg/ddac149.

Hirano, Michio, Alexander Shtilbans, Richard Mayeux, Mercy M. Davidson, Salvatore DiMauro, James A. Knowles, and Eric A. Schon. 1997. “Apparent MtDNA Heteroplasmy in Alzheimer’s Disease Patients and in Normals Due to PCR Amplification of Nucleus-Embedded MtDNA Pseudogenes.” Proc Natl Acad Sci U S A 94 (26): 14894–99. https://doi.org/10.1073/pnas.94.26.14894.

Khrapko, K., P. C. Andre, R. Cha, G. Hu, and W. G. Thilly. 1994. “Mutational Spectrometry: Means and Ends.” Progress in Nucleic Acid Research and Molecular Biology 49: 285–312.

Lander, E. S., and H. Lodish. 1990. “Mitochondrial Diseases: Gene Mapping and Gene Therapy.” Cell 61 (6): 925–26.

Lutz-Bonengel, Sabine, and Walther Parson. 2019. “No Further Evidence for Paternal Leakage of Mitochondrial DNA in Humans Yet.” Proceedings of the National Academy of Sciences 116 (6): 1821–22. https://doi.org/10.1073/pnas.1820533116.

Otten, Auke B. C., Suzanne C. E. H. Sallevelt, Phillippa J. Carling, Joseph C. F. M. Dreesen, Marion Drüsedau, Sabine Spierts, Aimee D. C. Paulussen, et al. 2018. “Mutation-Specific Effects in Germline Transmission of Pathogenic MtDNA Variants.” Hum Reprod 33 (7): 1331–41.

Popadin, Konstantin, Konstantin Gunbin, Leonid Peshkin, Sofia Annis, Zoe Fleischmann, Melissa Franco, Yevgenya Kraytsberg, Natalya Markuzon, Rebecca R. Ackermann, and Konstantin Khrapko. 2022. “Mitochondrial Pseudogenes Suggest Repeated Inter-Species Hybridization among Direct Human Ancestors.” Genes 13 (5): 810. https://doi.org/10.3390/genes13050810.

Popadin, Konstantin, Leonard V. Polishchuk, Leila Mamirova, Dmitry Knorre, and Konstantin Gunbin. 2007. “Accumulation of Slightly Deleterious Mutations in Mitochondrial Protein-Coding Genes of Large versus Small Mammals.” Proceedings of the National Academy of Sciences 104 (33): 13390–95.

Stewart, J. B., C. Freyer, J. L. Elson, A. Wredenberg, Z. Cansu, A. Trifunovic, and N. G. Larsson. 2008. “Strong Purifying Selection in Transmission of Mammalian Mitochondrial DNA.” PLoS Biol 6 (1): e10.

Wallace, Douglas C., Carol Stugard, Deborah Murdock, Theodore Schurr, and Michael D. Brown. 1997. “Ancient MtDNA Sequences in the Human Nuclear Genome: A Potential Source of Errors in Identifying Pathogenic Mutations.” Proceedings of the National Academy of Sciences 94 (26): 14900–905. https://doi.org/10.1073/pnas.94.26.14900.

Wei, Wei, Alistair T. Pagnamenta, Nicholas Gleadall, Alba Sanchis-Juan, Jonathan Stephens, John Broxholme, Salih Tuna, et al. 2020. “Nuclear-Mitochondrial DNA Segments Resemble Paternally Inherited Mitochondrial DNA in Humans.” Nature Communications 11 (1): 1740. https://doi.org/10.1038/s41467-020-15336-3.

Wei, Wei, Salih Tuna, Michael J. Keogh, Katherine R. Smith, Timothy J. Aitman, Phil L. Beales, David L. Bennett, et al. 2019. “Germline Selection Shapes Human Mitochondrial DNA Diversity.” Science 364 (6442): eaau6520.

Zhang, Haixin, Marco Esposito, Mikael G. Pezet, Juvid Aryaman, Wei Wei, Florian Klimm, Claudia Calabrese, et al. 2021. “Mitochondrial DNA Heteroplasmy Is Modulated during Oocyte Development Propagating Mutation Transmission.” Science Advances 7 (50): eabi5657. https://doi.org/10.1126/sciadv.abi5657.

