## Supplement for "Reanalysis of mtDNA mutations of human primordial germ cells (PGCs) reveals NUMT contamination and suggests that selection in PGCs may be positive"

### Fleischmann et al., Supplement

#### Table of contents:


| | Page: |
|---|---|
| Supplementary Table | 2 |
| Supplementary Figure S1 | 3 |
| Supplementary Notes: | 5 |
| Note 1 | 5 |
| Note 2 | 6 |
| Note 3 | 6 |
| Note 4 | 7 |

### Supplementary Table

| Sample | Position | Ref | Alt | NuMT | Syn | Rep | MAF |
|---|---|---|---|---|---|---|---|
| **Early PGCs** | | | | | | | |
| CS12-12108 | 8642 | A | G | | NSyn | 1 | 0.1045 |
| CS12-12108 | 14981 | A | G | | NSyn | 1 | 0.1009 |
| CS12-12108 | 7034 | C | A | | NSyn | 9 | 0.0142 |
| CS12-12222 | 9918 | G | A | ch5 | NSyn | 9 | 0.016 |
| CS12-12108 | 7961 | T | C | | Syn | 11 | 0.0134 |
| CS12-12222 | 7805 | G | T | | NSyn | 12 | 0.0215 |
| CS12-12108 | 7805 | G | T | | NSyn | 12 | 0.0127 |
| CS12-12108 | 14390 | C | T | | NSyn | 12 | 0.0138 |
| **Late PGCs** | | | | | | | |
| CS21-11595 | 3918 | G | A | | Syn | 1 | 0.0147 |
| CS21-12061 | 5450 | C | T | | Syn | 1 | 0.025 |
| CS20-11593 | 8519 | G | A | | NSyn | 1 | 0.01 |
| CS20-11941 | 11641 | A | G | | Syn | 1 | 0.0112 |
| CS21-12005 | 7034 | C | A | | NSyn | 9 | 0.0112 |
| CS20-11593 | 7034 | C | A | | NSyn | 9 | 0.0105 |
| CS21-12064 | 7805 | G | T | | NSyn | 12 | 0.0201 |
| CS21-12061 | 7805 | G | T | | NSyn | 12 | 0.0189 |
| CS20-11593 | 7805 | G | T | | NSyn | 12 | 0.0164 |
| CS21-12005 | 7805 | G | T | | NSyn | 12 | 0.0122 |
| CS20-11941 | 7805 | G | T | | NSyn | 12 | 0.0121 |
| CS20-11593 | 7961 | T | C | | Syn | 11 | 0.0144 |
| CS21-12061 | 7961 | T | C | | Syn | 11 | 0.0142 |
| CS21-12064 | 7961 | T | C | | Syn | 11 | 0.0118 |
| CS20-11593 | 9380 | G | A | ch5 | Syn | 4 | 0.0101 |
| CS20-11593 | 9428 | T | C | ch5 | Syn | 5 | 0.0109 |
| CS20-11593 | 9429 | G | A | ch5 | NSyn | 5 | 0.0104 |
| CS20-11593 | 9506 | C | T | ch5 | Syn | 5 | 0.01 |
| CS20-11593 | 9540 | T | C | ch5 | Syn | 6 | 0.0124 |
| CS20-11593 | 9548 | G | A | ch5 | Syn | 7 | 0.0127 |
| CS20-11593 | 9554 | G | A | ch5 | Syn | 6 | 0.0113 |
| CS20-11593 | 9911 | C | T | ch5 | Syn | 8 | 0.0129 |
| CS20-11941 | 9911 | C | T | ch5 | Syn | 8 | 0.0106 |
| CS21-12005 | 9918 | G | A | ch5 | NSyn | 9 | 0.0101 |
| CS21-12064 | 14390 | C | T | | NSyn | 12 | 0.0188 |
| CS21-12061 | 14390 | C | T | | NSyn | 12 | 0.0151 |
| CS20-11593 | 14390 | C | T | | NSyn | 12 | 0.0136 |
| CS21-12005 | 14390 | C | T | | NSyn | 12 | 0.012 |

**Footnote:**

**NuMT**: '**ch5**' identifies confirmed SNV in the nuclear genome, chromosome

**Syn**: Synonymity of the SNV. Synonymous (Syn) or nonsynonymous (NSyn)

**Rep**: Repetition, i.e., number of samples (of 12 total) where SNV has exceeded 0.3%

**MAF**: Minor Allele Frequency. In this context, an equivalent of mutant fraction.

Light blue shading: the most highly NuMT-contaminated sample, CS20-11593

Light orange shading: repeated mutations

Purple shading: high fraction mutations likely originating from previous generation

### Supplementary Figure S1

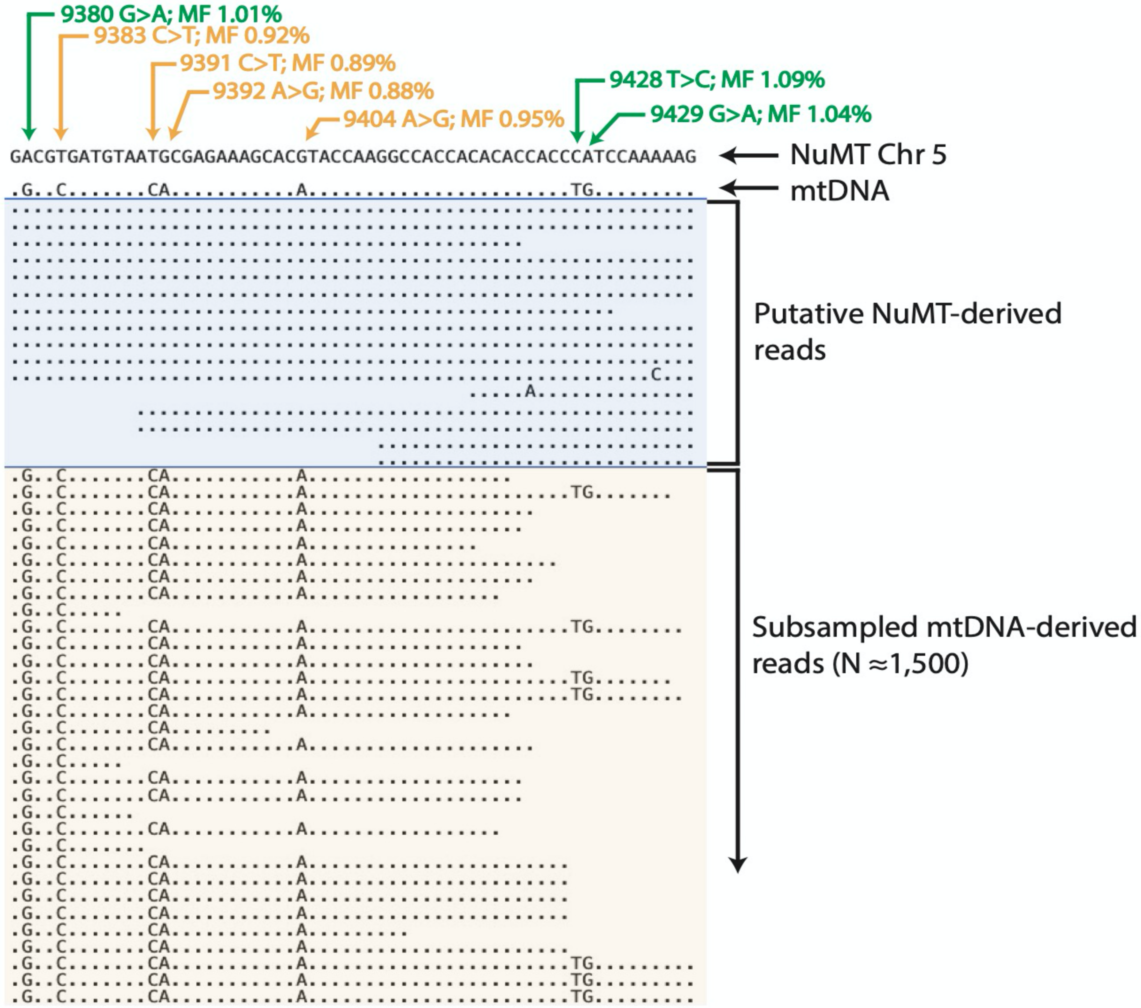


**Figure S1. A substantial proportion (about 1%) of next generation primary sequencing reads in the Floros et al. data are NUMT-derived reads identified by the presence of *multiple* NUMT-specific sequence variants that all map to the same NUMT sequence on chromosome 5.**

Shown here is a representative section of an alignment of the next generation sequencing reads from the most NUMT-contaminated late PGC sample (CS20-11593) aligned to the genomic sequence of the NUMT located on chromosome 5 (hg38(100045938-100055045), label: 'NUMT Chr 5') and to the human mitochondrial genome (label: 'mtDNA'). The variants are identified by their position in the mitochondrial genome and their reported mutant fractions are shown in yellow and green labels at the top. The dots represent bases identical to the NUMT reference. In this section of the alignment, NUMT reads are identified by 7 SNVs mapping to the NUMT sequence (the corresponding nucleotides are shown in mtDNA reads, not in NUMT reads, because NUMT is the reference). **Green** marks SNVs that, according to Floros et al criterion, were 'approved' real mutations with fraction> 1%. **Yellow** are mutations wit MF<1% which were considered not real mutations.

We do not know what causes subtle variation of the ratio of NUMT-to-mtDNA reads from SNV to SNV. But we do see that because fraction of NUMT reads in this sample is so close to the 1% threshold, even very small variations are sufficient to move an SNV from approved to not approved status. Changes so small could result from random variations of the number of reads. More important question is why this particular late pooled PGC sample CS20-11593 became far more contaminated with NUMT SNVs than other samples (8 of 11 NUMT SNVs are from CS20-11593) and thereby almost single-handedly caused the false perception of purifying selection. DNA of the samples is not available for re-analysis, so we may never know what exactly happened in these experiments, but a plausible hypothesis can be offered. We note that differential PCR-blocking damage of the mtDNA vs. nuclear DNA will result in different proportions of mtDNA- and nDNA-derived PCR product. According to our observations (Kraytsberg et al. 2009) 10.1007/978-1-59745-521-3_21, some copies of mtDNA (but less so nDNA) are rendered unamplifiable, apparently due to damage. If mtDNA damage in CS20-11593 was relatively slightly higher than in other samples, that would explain higher prevalence of NUMT contamination of this sample. Differences in damage could have resulted from subtle differences in the handling of the sample.

*Note on figure construction*: To be able to fit sufficient data into the figure, the reads were subsampled so that the total coverage of this region of the mitochondrial genome was about 1,500. The number of NUMT-mapping reads in the alignment is 15, i.e., ~1% of total reads, which is close to the mutant fraction of NUMT-mapping mutations in this sample and explains why mutant fractions of SNVs tend to 'hoover' around 1%., which results in variable 'endorsement' of SNVs. This disrupted the anticipated 'block' appearance of NUMT-derived SNVs and apparently made their recognition as such difficult.

### Supplementary Notes.

**Supplementary Note 1: Additional analysis of NuMT co-amplification in the pooled PGC sample.**

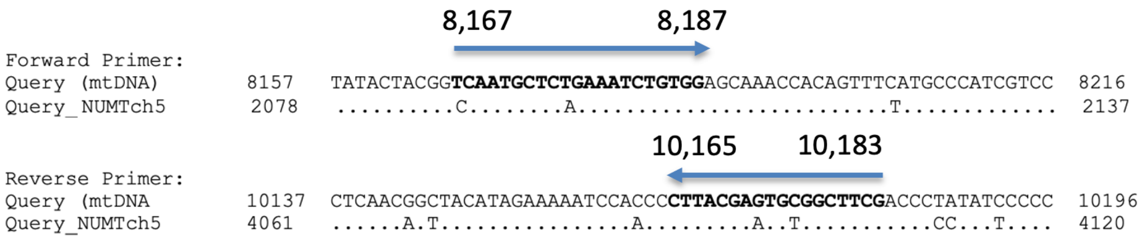


#### Primer pair that co-amplifies NUMT ch5

Scheme above shows the position and homology to NUMT ch5 of the primer set that caused co-amplification of NUMT ch5 in the pooled PGC samples. The primers' sequences are shown as aligned to the mtDNA genome, and the corresponding priming sites in the NUMT sequence show only the differences (matches are depicted as dots). Note how well the primers and the NUMT sequence match (there are only two mismatches per primer far from the mismatch-sensitive 3' end of the primer).

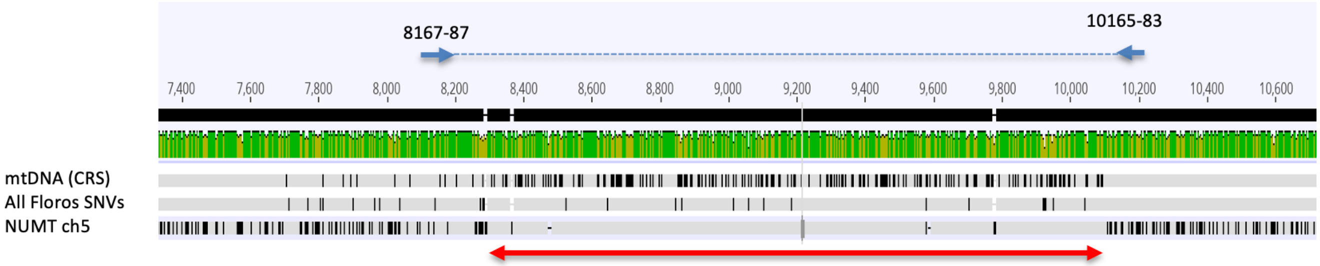


Shown above is the **"SNV footprint"** of the NUMT, i.e., area of the mitochondrial genome where almost all SNVs detected by the Floros et al. study is derived from the NUMT on chr5 (the position of the 'footprint' is shown by the red double arrow). **Indeed,** the striking *scarcity of vertical black stripes in the part* of the NUMT sequence (bottom sequence) marked by the double arrow means that SNVs in that area are identical to Floros SNVs (middle sequence). As expected, the footprint is flanked by PCR primers used to amplify this region of mtDNA which confirms that that NUMT was co-amplified by PCR with this pair of primers.

**Methodology**: This is an alignment of NUMT ch5 sequence (used as reference in this alignment), the mtDNA sequence mtDNA (CRS – Cambridge Revised Sequence), and an artificial sequence ("All Floros SNVs"). The latter was made by adding all SNVs reported in pooled PGC dataset of Floros et al to the mtDNA sequence. This includes *all* mutations reported, i.e., including those below the 1% threshold, down to 0.3%. **Explanation**: Black vertical lines indicate differences with the consensus sequence. Normally these differences are expected to be seen in the NUMT sequence, which is about 5% divergent from mtDNA. This is what is seen to the right and to the left of the PCR fragment. However, within the PCR fragment, the distribution of differences 'switches paths' and they all now appear in the mtDNA, because consensus now is between 'All Floros SNVs' and 'NUMTch5'. This demonstrates that a great majority of SNVs from pooled PGC detected in this region originate in NUMT. Indeed, a black stripe at an SNV's position in mtDNA means that corresponding SNV is identical in the NUMT and the 'All Floros SNVs' sequence. The 'SNV footprint' thus is a strong confirmation of NUMT co-amplification with this pair of primers. Note that only a portion of co-amplified SNVs made it to the 'endorsed mutations' set, because most of them failed to reach 1% threshold, as explained in the caption of FigS1.

**Supplementary Note 2: Single PGC PCR primers do not amplify NUMT Ch5**

The map below demonstrates that neither of the two primer pairs used to amplify mtDNA from single PGCs are able to co-amplify NUMT ch5 because one of the primers in each of the pairs is positioned outside of the NUMT sequence.

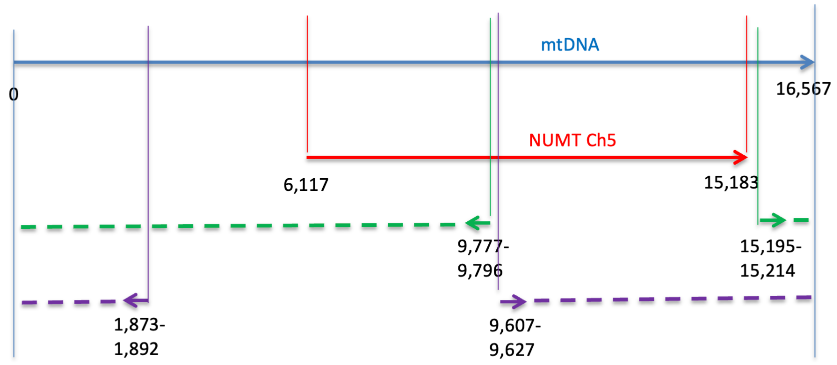


Single PGC primer sets

**Supplementary Note 3: NuMT-derived SNVs are highly synonymous.**

Perhaps counterintuitively, NuMT-derived mutations (i.e., mutations of nuclear pseudogenes of mtDNA) are overall highly synonymous, in contrast to highly nonsynonymous mutations in conventional nuclear pseudogenes. We have demonstrated this peculiar feature of NUMTs in humans (Popadin 2022), and this has been demonstrated previously by others for bristletails. NuMT variants are highly synonymous because a majority of NuMT variants are in fact ‘reverse mtDNA polymorphisms’. NuMT sequences, which reside in the protective nuclear DNA environment, mutate at a much lower rate than the mitochondrial genome, meaning that most of the differences between NuMTs and actual mtDNA sequences are mutations in mtDNA that were fixed in the species since the insertion of the NuMT into the nuclear genome. Thus, a majority of NuMT/mtDNA differences are historic mtDNA mutations, which are highly synonymous because most of them are fixed in the population by random drift and thus need to be fairly neutral to keep selection against them at a permissively low level.

**Supplementary Note 4:**

| | MF/ synony-mity | p-value |
|---|---|---|
| 1 | 0.4746 | 0.712 |
| 2 | 0.3721 | 0.505 |
| 3 | 0.3167 | 0.358 |
| 4 | 0.2953 | 0.253 |
| 5 | 0.1419 | 0.179 |
| 6 | 0.1286 | 0.125 |
| 7 | 0.1241 | 0.088 |
| 8 | 0.1229 | 0.062 |
| 9 | 0.1011 | 0.044 |
| 10 | 0.0962 | 0.030 |
| 11 | 0.0927 | 0.021 |
| 12 | 0.0906 | 0.015 |
| 13 | 0.0826 | 0.015 |
| 14 | 0.0824 | 0.014 |
| 15 | 0.0812 | 0.012 |
| 16 | 0.0802 | 0.024 |
| 17 | 0.0783 | 0.019 |
| 18 | 0.0744 | 0.016 |
| 19 | 0.0738 | 0.013 |
| 20 | 0.0735 | 0.010 |
| 21 | 0.0726 | 0.021 |
| 22 | 0.0723 | 0.017 |
| 23 | 0.0711 | 0.014 |
| 24 | 0.0691 | 0.011 |
| 25 | 0.0682 | 0.009 |
| 26 | 0.0665 | 0.017 |
| 27 | 0.0643 | 0.014 |
| 28 | 0.0601 | 0.023 |
| 29 | 0.0593 | 0.020 |
| 30 | 0.0589 | 0.030 |
| 31 | 0.0579 | 0.026 |
| 32 | 0.0566 | 0.038 |
| 33 | 0.055 | 0.034 |
| 34 | 0.0536 | 0.029 |
| 35 | 0.0516 | 0.026 |
| 36 | 0.0505 | 0.036 |
| 37 | 0.0477 | 0.032 |
| 38 | 0.0454 | 0.043 |
| 39 | 0.0452 | 0.038 |
| 40 | 0.0447 | 0.050 |
| 41 | 0.0434 | 0.063 |
| 42 | 0.0427 | 0.078 |
| 43 | 0.0424 | 0.072 |
| 44 | 0.0418 | 0.066 |
| 45 | 0.0406 | 0.060 |
| 46 | 0.0405 | 0.074 |
| 47 | 0.0403 | 0.069 |
| 48 | 0.0402 | 0.083 |
| 49 | 0.0396 | 0.077 |
| 50 | 0.0384 | 0.072 |

**Decrease of synonymity at high mutant fractions in single PGC mutations is statistically significant.**

As discussed in main text, purifying selection in PGCs predicts an increased synonymity of high fraction single PGC mutations. Equally, it predicts that non-synonymous mutations on average should be of lower mutant fraction than synonymous ones. In contradiction to the latter prediction, average mutant fraction of synonymous single PGC mutations was significantly lower than that of non-synonymous mutations (0.028 vs. 0.042, $p<0.02$, two sample t-test; primary data shown below in the 'Data' section).

Interestingly, a finer statistical analysis(www.statskingdom.com) revealed that the single PGC data is highly non-normal (p<0.00001, Shapiro Wilk test) and contained a subset of 15 outliers (Tukey's fences, k=1.5), i.e., a group of 14 high-mutant fraction mutations among non-synonymous mutations, and only 1 outlier among synonymous mutations. Among outliers, there are 12 non-synonymous mutations exceeding any synonymous mutation in the data set by the mutant fraction (See **table on the left**, which shows mutant fractions and synonymity (**red** – non-synonymous, **green** – synonymous) of the top 50 protein coding single PGC mutations, ranked by mutant fraction). Note a visually apparent enrichment of nonsynonymous mutations among the top (high mutant fraction) mutations – the "red cluster".

As noted in the main text, the very existence of a high mutant fraction 'cluster' of nonsynonymous mutations contradicts purifying selection. We therefore estimated statistical significance of the observation of this 'cluster'. Under no-selection scenario (null hypothesis) we can use binomial probability to estimate the likelihood of observing 12 nonsynonymous mutations in a row among highest MAF mutations. The *a prori* binomial probability of a group of 12 nonsynonymous mutations in the top MF under null hypotheses of full randomness is 0.71^12 = 0.017; (where 0.71 is the fraction of nonsynonymous mutation among protein coding mutations in the entire dataset). Hence the existence of this group is a significant finding at least with p~0.017.

We further confirmed and generalized this conservative binomial estimate by a bootstrap simulation described in the "Methodology" paragraph below. In this simulation, we tested whether apparent enrichment of nonsynonymous mutations at high mutant fractions (see Table on the left) is a significant observation. We tested significance of enrichment for all possible 'top MAF subsets' of mutations, and, as shown in the table on the right, were able to robustly demonstrate significance of enrichment in a wide range of subsets: from subsets with MAF>=0.045 to those with MAF>=0.1011 (blue cells in the p-value column). p-values stay robustly below 0.05 and even occasionally descend below 0.01, indicating high significance (dark blue). This robustness excludes a possibility that the observed effect is due to the 'cherry-picking' bias (i.e., particular combination of parameters leading to spurious significance). Note that p-value stays above 0.05 in the top 8 rows of the table: this is merely because the number of datapoints is too low to ensure significance (this is in accord with the higher binomial probability of smaller sets).

Because enrichment of nonsynonymous mutations towards higher MAFs is significant and because this trend is incompatible with purifying section in PGCs, we conclude that the hypothesis of purifying (negative) selection is rejected.

**Methodology**:
For bootstrap simulation, we emulated the null hypothesis (i.e. lack relationship between synonymity and mutant fraction) by randomizing the original list of mutant fraction/synonymity data pairs of protein coding single PGC mutations (MF>=1% - 188 datapoints). We repeated 100,000-bootstrap simulation twice and confirmed reproducibility of the p-value estimates to the third digit. For the original data set and

for each randomization, we sorted the data by mutant fraction and calculated weighted average synonymity for the set of progressively increasing subsets of top-MAF mutations starting from the top of the list. First window included the highest MAF mutation (MAF=0.4746), second subset - two highest MAF mutations (0.4746 and 0.3721), …, and so on till the last widow that included all 188 mutations. Then, for each subset, we counted the number of bootstraps where weighted average synonymity was equal or lower than that observed in the corresponding subset of original data set. The fraction of these bootstraps was considered the non-parametric p-value for the particular set of top-MAF mutations. Weighted synonymity was calculated by adding mutant fractions of all synonymous mutations in the subset, subtracting mutant fractions of all non-synonymous mutations in the subset and dividing the result by the sum of all mutant fractions in the subset. Thus, weighted synonymity changes from -1 for a set of fully nonsynonymous mutations to +1 for a set of fully synonymous mutations and is zero when total mutant fraction of synonymous mutations in a set is equal to that of nonsynonymous mutations.

**Data (**single PGC, protein coding mutations, MF>1%, derived from (Floros 2022), ranked by mutant fraction).

| | Syn / Non-Syn | MF |
|---|---|---|
| 1 | nonsynonymous | 0.4746 |
| 2 | nonsynonymous | 0.3721 |
| 3 | nonsynonymous | 0.3167 |
| 4 | nonsynonymous | 0.2953 |
| 5 | nonsynonymous | 0.1419 |
| 6 | nonsynonymous | 0.1286 |
| 7 | nonsynonymous | 0.1241 |
| 8 | nonsynonymous | 0.1229 |
| 9 | nonsynonymous | 0.1011 |
| 10 | nonsynonymous | 0.0962 |
| 11 | stopgain | 0.0927 |
| 12 | nonsynonymous | 0.0906 |
| 13 | synonymous | 0.0826 |
| 14 | nonsynonymous | 0.0824 |
| 15 | nonsynonymous | 0.0812 |
| 16 | synonymous | 0.0802 |
| 17 | nonsynonymous | 0.0783 |
| 18 | nonsynonymous | 0.0744 |
| 19 | nonsynonymous | 0.0738 |
| 20 | nonsynonymous | 0.0735 |
| 21 | synonymous | 0.0726 |
| 22 | nonsynonymous | 0.0723 |
| 23 | nonsynonymous | 0.0711 |
| 24 | nonsynonymous | 0.0691 |
| 25 | stopgain | 0.0682 |
| 26 | synonymous | 0.0665 |
| 27 | nonsynonymous | 0.0643 |
| 28 | synonymous | 0.0601 |
| 29 | nonsynonymous | 0.0593 |
| 30 | synonymous | 0.0589 |
| 31 | nonsynonymous | 0.0579 |

| | | |
|---|---|---|
| 32 | synonymous | 0.0566 |
| 33 | nonsynonymous | 0.055 |
| 34 | nonsynonymous | 0.0536 |
| 35 | nonsynonymous | 0.0516 |
| 36 | synonymous | 0.0505 |
| 37 | nonsynonymous | 0.0477 |
| 38 | synonymous | 0.0454 |
| 39 | stopgain | 0.0452 |
| 40 | synonymous | 0.0447 |
| 41 | synonymous | 0.0434 |
| 42 | synonymous | 0.0427 |
| 43 | nonsynonymous | 0.0424 |
| 44 | nonsynonymous | 0.0418 |
| 45 | nonsynonymous | 0.0406 |
| 46 | synonymous | 0.0405 |
| 47 | nonsynonymous | 0.0403 |
| 48 | synonymous | 0.0402 |
| 49 | stopgain | 0.0396 |
| 50 | nonsynonymous | 0.0384 |
| 51 | nonsynonymous | 0.038 |
| 52 | synonymous | 0.0362 |
| 53 | synonymous | 0.0351 |
| 54 | nonsynonymous | 0.0344 |
| 55 | nonsynonymous | 0.0334 |
| 56 | nonsynonymous | 0.0329 |
| 57 | nonsynonymous | 0.0325 |
| 58 | nonsynonymous | 0.0324 |
| 59 | synonymous | 0.0316 |
| 60 | synonymous | 0.0312 |
| 61 | nonsynonymous | 0.0308 |
| 62 | nonsynonymous | 0.0306 |
| 63 | nonsynonymous | 0.0305 |
| 64 | stopgain | 0.0304 |
| 65 | nonsynonymous | 0.0296 |
| 66 | nonsynonymous | 0.0294 |
| 67 | nonsynonymous | 0.0289 |
| 68 | synonymous | 0.0287 |
| 69 | nonsynonymous | 0.0284 |
| 70 | nonsynonymous | 0.0282 |
| 71 | synonymous | 0.0281 |
| 72 | nonsynonymous | 0.0272 |
| 73 | nonsynonymous | 0.0267 |
| 74 | nonsynonymous | 0.0264 |

| | | |
|---|---|---|
| 75 | synonymous | 0.0263 |
| 76 | synonymous | 0.0263 |
| 77 | nonsynonymous | 0.0261 |
| 78 | stopgain | 0.0257 |
| 79 | nonsynonymous | 0.0248 |
| 80 | nonsynonymous | 0.0248 |
| 81 | synonymous | 0.0247 |
| 82 | synonymous | 0.0243 |
| 83 | nonsynonymous | 0.0242 |
| 84 | synonymous | 0.0236 |
| 85 | synonymous | 0.0235 |
| 86 | synonymous | 0.0233 |
| 87 | nonsynonymous | 0.0232 |
| 88 | nonsynonymous | 0.0232 |
| 89 | nonsynonymous | 0.0222 |
| 90 | nonsynonymous | 0.0219 |
| 91 | nonsynonymous | 0.0218 |
| 92 | nonsynonymous | 0.0209 |
| 93 | nonsynonymous | 0.0207 |
| 94 | synonymous | 0.0201 |
| 95 | nonsynonymous | 0.0199 |
| 96 | synonymous | 0.0196 |
| 97 | nonsynonymous | 0.0194 |
| 98 | nonsynonymous | 0.0194 |
| 99 | nonsynonymous | 0.0192 |
| 100 | nonsynonymous | 0.0189 |
| 101 | nonsynonymous | 0.0189 |
| 102 | nonsynonymous | 0.0187 |
| 103 | nonsynonymous | 0.0183 |
| 104 | nonsynonymous | 0.0183 |
| 105 | nonsynonymous | 0.0183 |
| 106 | nonsynonymous | 0.0182 |
| 107 | nonsynonymous | 0.0175 |
| 108 | nonsynonymous | 0.0174 |
| 109 | nonsynonymous | 0.0174 |
| 110 | synonymous | 0.0173 |
| 111 | nonsynonymous | 0.0172 |
| 112 | nonsynonymous | 0.0172 |
| 113 | nonsynonymous | 0.0172 |
| 114 | synonymous | 0.0172 |
| 115 | nonsynonymous | 0.0169 |
| 116 | synonymous | 0.0167 |
| 117 | nonsynonymous | 0.0166 |

| | | |
|---|---|---|
| 118 | nonsynonymous | 0.0165 |
| 119 | nonsynonymous | 0.0165 |
| 120 | nonsynonymous | 0.0162 |
| 121 | synonymous | 0.0162 |
| 122 | synonymous | 0.0161 |
| 123 | synonymous | 0.0161 |
| 124 | nonsynonymous | 0.0158 |
| 125 | nonsynonymous | 0.0155 |
| 126 | nonsynonymous | 0.0155 |
| 127 | nonsynonymous | 0.0155 |
| 128 | nonsynonymous | 0.0154 |
| 129 | nonsynonymous | 0.0151 |
| 130 | nonsynonymous | 0.0151 |
| 131 | nonsynonymous | 0.015 |
| 132 | synonymous | 0.0149 |
| 133 | nonsynonymous | 0.0148 |
| 134 | nonsynonymous | 0.0146 |
| 135 | nonsynonymous | 0.0145 |
| 136 | nonsynonymous | 0.0145 |
| 137 | synonymous | 0.0143 |
| 138 | synonymous | 0.0141 |
| 139 | nonsynonymous | 0.0138 |
| 140 | synonymous | 0.0135 |
| 141 | synonymous | 0.0133 |
| 142 | synonymous | 0.0132 |
| 143 | nonsynonymous | 0.0131 |
| 144 | nonsynonymous | 0.0131 |
| 145 | stopgain | 0.0131 |
| 146 | synonymous | 0.0131 |
| 147 | nonsynonymous | 0.013 |
| 148 | nonsynonymous | 0.0129 |
| 149 | nonsynonymous | 0.0127 |
| 150 | nonsynonymous | 0.0126 |
| 151 | nonsynonymous | 0.0125 |
| 152 | nonsynonymous | 0.0123 |
| 153 | stopgain | 0.0123 |
| 154 | nonsynonymous | 0.0122 |
| 155 | nonsynonymous | 0.0121 |
| 156 | nonsynonymous | 0.012 |
| 157 | synonymous | 0.0119 |
| 158 | synonymous | 0.0117 |
| 159 | nonsynonymous | 0.0115 |
| 160 | nonsynonymous | 0.0114 |

| | | |
|---|---|---|
| 161 | synonymous | 0.0114 |
| 162 | synonymous | 0.0114 |
| 163 | synonymous | 0.0113 |
| 164 | nonsynonymous | 0.0112 |
| 165 | nonsynonymous | 0.0111 |
| 166 | nonsynonymous | 0.0111 |
| 167 | nonsynonymous | 0.0111 |
| 168 | nonsynonymous | 0.0111 |
| 169 | stopgain | 0.0111 |
| 170 | nonsynonymous | 0.0109 |
| 171 | nonsynonymous | 0.0108 |
| 172 | synonymous | 0.0108 |
| 173 | nonsynonymous | 0.0107 |
| 174 | nonsynonymous | 0.0106 |
| 175 | nonsynonymous | 0.0106 |
| 176 | nonsynonymous | 0.0106 |
| 177 | nonsynonymous | 0.0106 |
| 178 | nonsynonymous | 0.0104 |
| 179 | synonymous | 0.0104 |
| 180 | nonsynonymous | 0.0103 |
| 181 | nonsynonymous | 0.0103 |
| 182 | nonsynonymous | 0.0102 |
| 183 | nonsynonymous | 0.0102 |
| 184 | synonymous | 0.0102 |
| 185 | synonymous | 0.0102 |
| 186 | synonymous | 0.0101 |
| 187 | nonsynonymous | 0.01 |
| 188 | synonymous | 0.01 |
